# Divergent gene expression levels between diploid and autotetraploid *Tolmiea* (Saxifragaceae) relative to the total transcriptome, the cell, and biomass

**DOI:** 10.1101/169367

**Authors:** Clayton J. Visger, Gane K-S. Wong, Yong Zhang, Pamela S. Soltis, Douglas E. Soltis

**Affiliations:** Department of Biological Sciences, California State University Sacramento, Sacramento, CA 95819, USA; Department of Biological Sciences, University of Alberta, Edmonton AB, T6G 2E9, Canada; Department of Medicine, University of Alberta, Edmonton AB, T6G 2E1, Canada; Bejing Genomics Institute-Shenzhen, Beishan Industrial Zone, Yantian District, Shenzhen 518083, China; HuaHan Gene Co. Ltd., 7F JianAnShanHai Building, No. 8000, Shennan Road, Futian District, Shenzhen 518040, China; Florida Museum of Natural History, University of Florida, Gainesville, FL 32611, USA; Genetics Institute, University of Florida, Gainesville, FL 32610, USA; Department of Biology, University of Florida, Gainesville, FL 32611, USA

**Author notes:** Corresponding Author: Clayton J. Visger.

## Abstract

- Studies of gene expression and polyploidy are typically restricted to characterizing differences in transcript concentration. Integrating multiple methods of transcript analysis, we document a difference in transcriptome size, and make multiple comparisons of transcript abundance in diploid and autotetraploid *Tolmiea*.
- We use RNA spike-in standards to identify and correct for differences in transcriptome size, and compare levels of gene expression across multiple scales: per transcriptome, per cell, and per biomass.
- In total, ~17% of all loci were identified as differentially expressed (DEGs) between the diploid and autopolyploid species. A shift in total transcriptome size resulted in only ~58% of the total DEGs being identified as differentially expressed following a per transcriptome normalization. When transcript abundance was normalized per cell, ~82% of the total DEGs were recovered. The discrepancy between per-transcriptome and per-cell recovery of DEGs occurs because per-transcriptome normalizations are concentration-based and therefore blind to differences in transcriptome size.
- While each normalization enables valid comparisons at biologically relevant scales, a holistic comparison of multiple normalizations provides additional explanatory power not available from any single approach. Notably, autotetraploid loci tend to conserve diploid-like transcript abundance per biomass through increased gene expression per cell, and these loci are enriched for photosynthesis-related functions.

## Introduction

Polyploidy (whole-genome duplication; WGD), a process now recognized to be of major importance across many eukaryotic lineages (Van de Peer et al., 2009; Jiao et al., 2011; Van de Peer, 2011; Jiao and Paterson, 2014), was long considered an evolutionary dead-end by some, including several of the most prominent evolutionary biologists of the past century (Stebbins, 1950; Wagner, 1970). However, during the last several decades there has been a resurgence of interest in the study of polyploid evolution, particularly the genetic and genomic consequences of polyploidy (e.g., Ainouche *et al*., 2012; Barker *et al*., 2016; Canestro, 2012; Chen and Birchler, 2013; Doyle, 2012; Gaeta *et al*., 2007; Doyle *et al*., 2008; Soltis and Soltis, 2009, 2012; Salmon *et al*., 2010; Greilhuber *et al*., 2012; Shi *et al*., 2012; Madlung and Wendel, 2013; Renny-Byfield and Wendel, 2014; Soltis *et al*., 2014; Wendel, 2015; Spoelhof *et al., accepted*).

The role of polyploidy in facilitating changes in gene expression, through expression level divergence, altered expression patterns (e.g., across tissue-types), and/or the generation of unique splice variants, is arguably one of the most important research topics in the field today (e.g., Liu *et al*., 2001; Adams *et al*., 2003; Adams and Wendel, 2012; Chelaifa *et al*., 2010; Dong and Adams, 2011; Ainouche *et al*., 2012; Buggs, 2012; Buggs *et al*., 2014; Rambani *et al*., 2014; see Yoo *et al*., 2014 for review). For example, autotetraploid *Solanum phureja* expresses ~10% of all loci at a different levels relative to diploid *S. phureaja* (Stupar et al., 2007). In *Paulownia fortunei*, the autotetraploid differentially expresses ~6% of all loci relative to the diploid progenitor (Zhang et al., 2014), and in *Arabidopsis thaliana* ~4% of all loci are differentially expressed between the diploid and autotetraploid derivative (Del Pozo and Ramirez-Parra, 2014). However, the rapid increase in studies of patterns of gene expression in polyploids appear to have outpaced our fundamental understanding of the transcriptome’s response to polyploidy *per se* and how this response might influence our interpretation of gene expression measures.

One major concern in comparisons of gene expression between diploids and polyploids is that any transcriptome amplification or transcriptome-wide effects induced following polyploidy are rarely, if ever, investigated (see Coate and Doyle, 2010, 2015). Transcriptional amplification is a biological phenomenon in which the total mRNA produced per cell is increased up to several fold in one treatment group compared to another, resulting in unequal total transcriptome sizes (Nie *et al*., 2012). Transcriptional amplification would be anticipated in a polyploid compared to a diploid progenitor, as polyploidy globally alters gene/genome copy number and often influences cell size, which, in turn, has been shown to correlate with transcriptome size (Fomina-yadlin *et al*., 2014).

Recent work has shed light on some of the biases inherent to expression level comparisons between treatment groups that differ in transcriptome size. Loven et al. (2012) showed that inferences drawn from a typical RNAseq workflow can be confounded when treatment groups have transcriptomes of differing size. Because transcriptome size variation has rarely been explored, RNAseq studies involving transcriptome amplification are likely to underestimate the proportion of the transcriptome being differentially expressed. Surprisingly, only a single investigation into gene expression change following polyploidy has accounted for variation in transcriptome size (Coate and Doyle, 2010). Coate and Doyle (2010) found that allotetraploid *G. dolicocarpa* has a total mRNA transcriptome ~1.4 times greater than its diploid progenitors, demonstrating that polyploidy can induce an increase in transcriptome size. However, similar data from additional polyploid systems are badly needed to achieve a broader understanding of the role that polyploidy plays in transcriptome-wide changes in gene expression. Current methods for comparing differences in levels of gene expression across the transcriptome rely on the detection of statistically different RNAseq read abundances at each locus to generate a summary of differential gene expression. RNAseq libraries are usually sequenced to different depths by chance, resulting in library size (the total number of reads per sample) varying across the final dataset. To prevent variation in library size from influencing analyses of differential expression, library size is typically normalized across all samples (e.g., Mortazavi *et al*., 2008). Commonly used library normalization methods quantify expression level on a concentration basis by dividing read abundance by a factor of the whole library (e.g., transcripts per million or reads per kilobase per million) (see Coate and Doyle, 2010, 2015; Lovén *et al*., 2012). Concentration-based normalizations place gene expression in the context of expression level per transcriptome. Using a concentration-based normalization approach to infer differences in expression level requires that total transcriptome size does not vary between ploidal levels. If transcriptome size varies by ploidy, then loci identified as differentially expressed (differentially expressed genes; DEGs) are not necessarily expressed at different levels, but instead are maintained in different concentrations relative to the transcriptome (Figure 1A).

**Figure 1:**
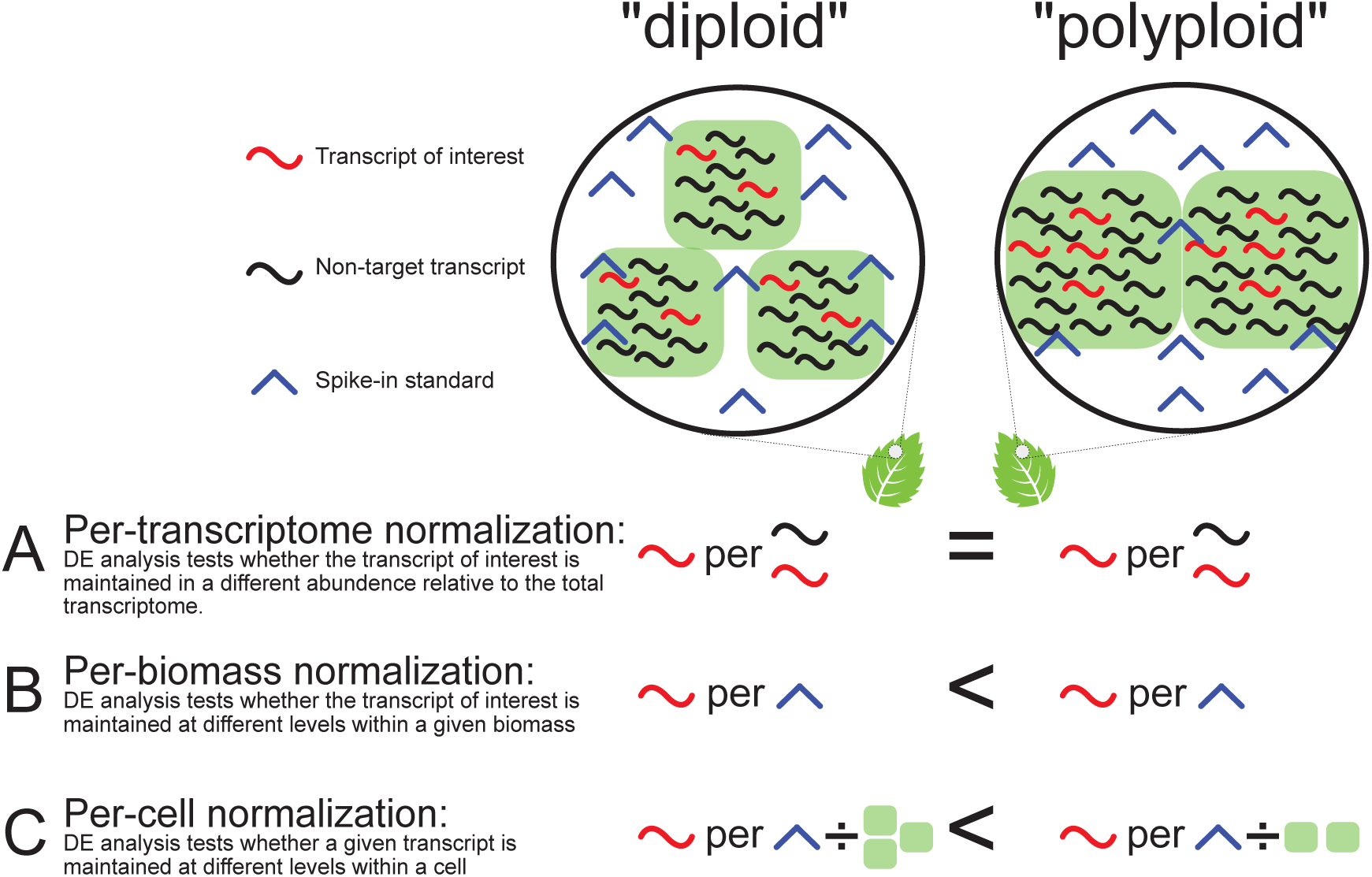
A simplified example of how spike-in standards can be used during read normalization to enable comparisons of expression level at different biological scales between a hypothetical diploid-polyploid pair with differing cell density. The large circles represent a unit of biomass and contain a number of cells (green squares). Beneath each circle is a depiction of how the read normalizations are calculated. Under a per transcriptome normalization, the ratio of target transcripts to the total transcriptome is compared. While the per biomass normalization uses the ratio of the transcript of interest to the spike-in transcripts. The per cell normalization also uses the ratio of the transcript of interest to spike-in transcripts, but scales the spike-in transcript abundance by cell density, represented here by multiplying the spike-in abundance by the number of contributing cells. Whether the transcript of interest would be found as not differentially expressed or higher/lower expressed in the polyploid under each normalization is indicted using ‘=’, ‘<’, or ‘>’ respectively.

To overcome these problems, the use of synthetic spike-in RNA standards has been proposed (Lovén *et al*., 2012). These standards, developed by the External RNA Controls Consortium (External RNA Controls Consortium, 2005; Baker *et al*., 2005; referred to as Spike-in RNA standards, henceforth spike-ins), facilitate comparison of absolute expression level across treatments with different transcriptome sizes (see Pine *et al*., 2016). Spike-ins allow for normalization of transcript abundance independent of transcriptome size. To date, spike-ins have been largely restricted to use in model systems (e.g., hamster – Fomina-yadlin *et al*., 2014; human – Xu *et al*., 2014; zebrafish – Schall *et al*., 2017), single cell sequencing (e.g., Krishnaswami *et al*., 2016; Liu *et al*., 2016), and methods development and validation (e.g., Lovén *et al*., 2012; Gu *et al*., 2014; Germain *et al*., 2016); they remain a promising but unutilized method for cross-ploidy comparisons.

Most polyploids exhibit differences in cell size relative to their parents, which may result in an alteration of cell density (Stebbins, 1971; Masterson, 1994). Because normalization per transcriptome is based on concentration of transcripts, these comparisons are therefore robust to variation in cell size and density. Differences in cell number and cell size might be expected to influence levels of gene expression on certain scales, but cell size and density are rarely investigated prior to studying differences in gene expression levels. Conversely, normalizing by the abundance of an internal standard within a library (e.g., spike-ins) does not account for variation in the number of contributing cells across treatments, and the inferred transcript abundance may be biased (Fomina-yadlin *et al*., 2014). It is therefore critical to have information on cell density differences between treatment groups when normalizing to an internal standard.

Following polyploidy, a balance of cell size and density effects, as well as differences in gene/allelic dosage, could have profound effects on all facets of plant physiology. Across a ploidal series in *Atriplex confertifolia*, cell density decreases with increasing ploidy (Warner and Edwards, 1989). Despite harboring fewer cells per unit area, higher ploidal levels of *A. confertifolia* are capable of higher photosynthesis per unit leaf area, due to increased photosynthesis of tetraploid cells relative to diploid cells. Conversely, in tetraploid *Medicago sativa*, which also exhibits lower cell density than does its diploid progenitor, photosynthesis per cell is increased in the tetraploid to yield diploid-like photosynthetic output per unit leaf area (Warner and Edwards, 1993). In *A. confertifolia* and *M. sativa*, cell size and density alone are not sufficient for predicting physiological change following polyploidy. Instead, information on the interaction between cell density and cell efficiency is needed to understand fully the physiological impact of polyploidy. In light of these observations, studies of changes in gene expression levels following polyploidy should routinely investigate the interaction between gene expression across a unit of biomass and per cell.

When discussing transcript abundance between samples of differing transcriptome sizes and/or differing cell density, a clear nomenclature is critical. We illustrate three separate ways of defining transcript abundance when both transcriptome size and cell density vary between treatment groups (Figure 1). When RNAseq data are normalized without an external standard (Figure 1A), differences in abundance observed between two treatments reflect a difference in transcript concentration. We follow Coate and Doyle (2010) in referring to this type of comparison as ‘per transcriptome’.

Differences in expression level between treatments normalized to the abundance of spike-ins reflect differences in transcript abundance relative to the abundance of spike-ins. When spike-ins are added in equal amounts to samples derived from equivalent volumes of tissue, then differences observed following a spike-in-based normalization indicate that transcripts differ in abundance within a given volume of tissue (Figure 1B). We refer to this comparison as ‘per biomass’. Finally, if the spike-in abundance is scaled by a factor equal to differences in cell density, differences in expression level will reflect changes in transcript abundance relative to the cell (Figure 1C), and we refer to this case as ‘per cell’ (Figure 1).

*Tolmiea* (Saxifragaceae) is an excellent evolutionary model for investigating the transcriptional impact of polyploidy because: 1) it is a clear diploid (*T. diplomenziesii*) and autotetraploid (*T. menziesii*) system; and 2) there is strong support for a single origin of the autotetraploid (Soltis and Soltis, 1988; Soltis *et al*., 1989; Visger *et al*., 2016). In addition, examination of changes in expression level following autopolyploidy is less problematic than similar investigations within allopolyploid systems for several reasons. First, autopolyploidy results in a duplication of a single genome, rather than the merger and duplication of two divergent genomes (as in allopolyploidy). Hence, a single diploid-based mapping reference can be used for both the diploid and the autotetraploid. Additionally, by definition, there are no homeologues in an autopolyploid, substantially reducing the potential for biased mismapping of paralogous reads. A single origin of the polyploid is also an advantage for exploring issues pertaining to transcriptional amplification and reduces analytical complexity. For example, multiple, independent origins of a polyploid may have differing effects on transcriptome size. Thus, an investigation accounting for biases in transcriptome size should start with an autopolyploid arising from a single origin.

Here we leverage spike-in RNA standards and multiple normalization methods to collectively characterize for the first-time gene expression across three biological scales in a natural system. Through this multi-scale comparison, we investigate how polyploidy has influenced transcriptional change in gene pathway stoichiometry, cellular expression levels, and transcript abundance across leaf organ tissue. Finally, we synthesize the differences represented across all three scales and place our findings within the context of ecophysiological data for *Tolmiea*.

## Materials and Methods

*Sampling*— Plants were collected from three geographically separate natural populations of both *T. diplomenziesii* and *T. menziesii* (Figure 2). Taking advantage of the ability of *Tolmiea* to reproduce asexually via plantlet formation, each individual collected in the field was subsequently propagated in quadruplicate in a greenhouse at the University of Florida. The resulting plantlets were then grown to maturity under standardized conditions within a common garden greenhouse at the University of Florida. One of four replicate plants from two *T. menziesii* populations died prior to sampling.

**Figure 2:**
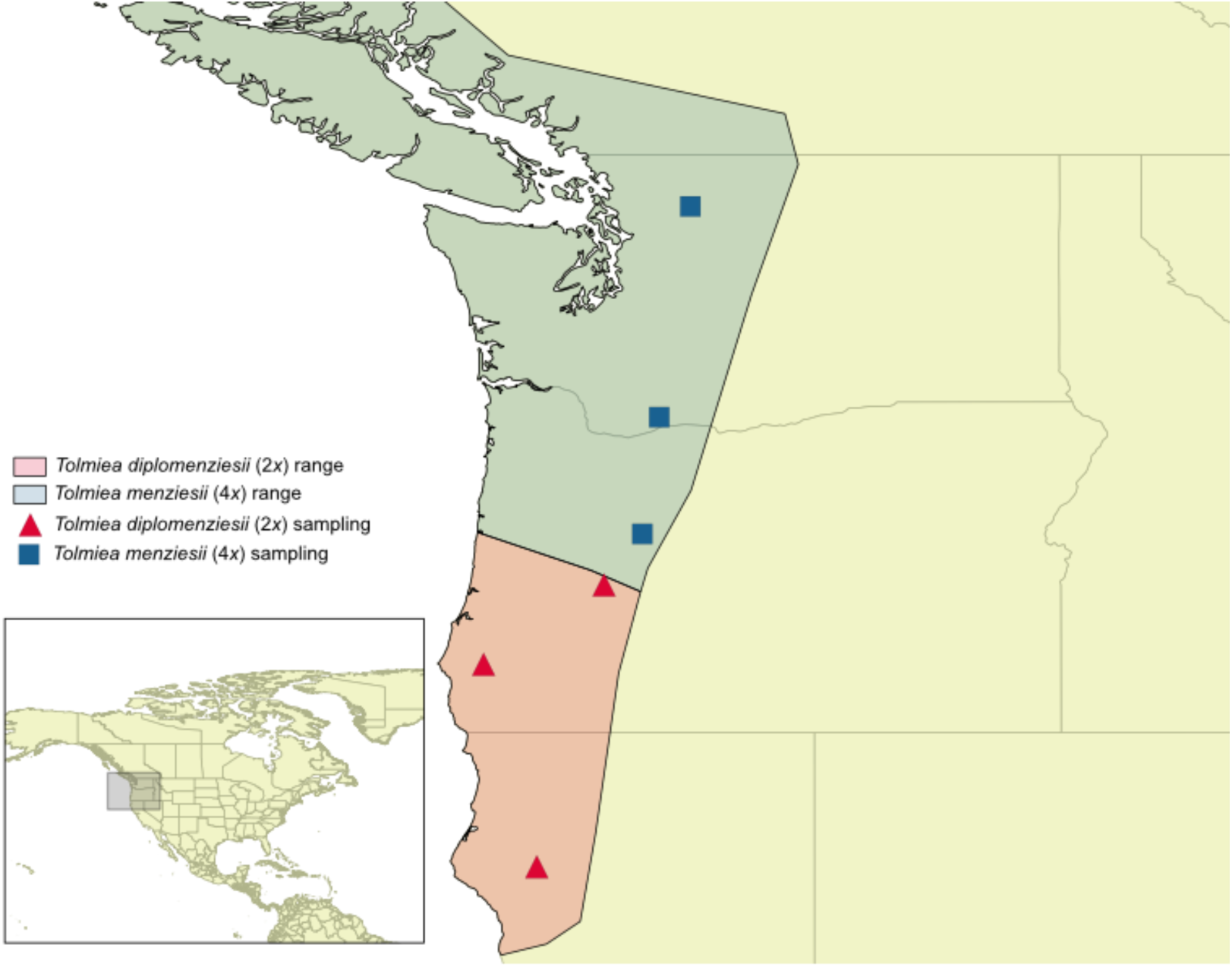
Generalized distributions of *Tolmiea menziesii* and *T. diplomenziesii*. The population sources for plants used in this study are represented as red triangles (*T. diplomenziesii*) and blue squares (*T. menziesii*).

Each of the 22 mature plants was sampled for equivalent volumes of leaf tissue using an 8-mm-diameter cork borer. The tissue was then flash frozen in liquid N2 and stored at -80C until RNA extraction. Total RNA was extracted from this tissue using the CTAB and Trizol method of Jordon-Thaden et al. (2015; protocol number 2) with the addition of 20% sarkosyl. DNA was removed using a Turbo DNA-free kit (Invitrogen, USA). Following the manufacturer’s recommendations, the total RNA was spiked with 4 ul of 1:100 diluted ERCC RNA Spike-in mix (Ambion, USA). RNAseq libraries were then built using the TruSeq kit (Illumina, USA); 100-bp paired-end sequencing was performed using an Illumina HiSeq at the Beijing Genomics Institute.

*Quantifying cell density*--To quantify differences in levels of gene expression per cell, it was necessary to account for differences in both transcriptome size and cell density between diploid and autotetraploid *Tolmiea*. A potential source of uncertainty is our estimation of differences in cell density that were used to normalize our reads on a relative per-cell basis. To decrease the likelihood of misrepresenting differences in cell density, we characterized cell density using both DNA/RNA co-extraction and cell counting as described below.

Duplicate leaf punches from each sample were used for co-extraction of DNA and RNA. We first followed the Jordon-Thaden et al. (2015) method #2 with 20% sakrosyl. Following the CTAB incubation, the supernatant was split into two equal aliquots, one of which was used for RNA extraction (following Jordon-Thaden et al., 2015) and the other for DNA extraction (following Doyle and Doyle, 1987). DNA concentrations were quantified with dsDNA broad-range chemicals using a Qubit (Life Technologies). DNA concentration was placed into a 1C context by dividing by ploidal level. The 1C DNA concentration was used to infer the relative difference in cell density of the leaf tissue contributing to RNA extraction between *T. menziesii* and *T. diplomenziesii*. To validate this approach, we also directly estimated cell density per unit area. Leaf punches two cm in diameter collected from 10 diploids and 11 tetraploids were digested in 500 ul of 10% chromic acid until cells were fully dissociated (Brown and Rickless, 1949; Ilut *et al*., 2012). Each individual was assayed twice, and the number of cells in the suspension was counted twice for each assay using 10-ul aliquots in a hemocytometer. Statistical analyses of 1C DNA concentration and cell density were performed using a linear mixed-effects model implemented in JMP (version 12; SAS Institute, Cary, NC, USA) with individual as a random effect. All datasets were tested for normality using a goodness of fit test, and if normality was rejected, the data were log transformed.

*Differential expression analysis*— Raw reads were cleaned using CutAdapt (Martin, 2011) and Sickle (Joshi and Fass, 2011). A *Tolmiea* reference transcriptome was generated from concatenated reads taken from all samples (with the spike-in reads removed), and *in silico* read normalization was employed using Trinity) (Grabherr *et al*., 2011). Extremely low-expressed isoforms were removed (< 1 transcript per million), and the remaining transcriptome was annotated using the Trinotate pipeline (http://trinotate.sourceforge.net/). Trimmed reads for each sample were mapped to a concatenation of the *Tolmiea* transcriptome with isoforms clustered together (using the Trinity ‘gene’ option) and the publically available ERCC spike-in reference using Bowtie2 (Langmead and Salzberg, 2012), and read counts were extracted using eXpress (Roberts, 2013).

We provide an iPython notebook including the entire code for the normalization, differential expression, and figure generation described below (Appendix 1). In short, read count normalization and differential expression analyses were conducted using Limma-Voom (Ritchie *et al*., 2015). A per-transcriptome normalization was implemented using the total library following the removal of spike-in count data to compute normalization factors. A per-biomass normalization used only spike-in count data to compute normalization factors. A per-cell normalization used the spike-in count data following *in silico* adjustment of tetraploid spike-in abundance using the difference of diploid versus tetraploid cell density (65%; see methods on quantifying cell size and density and Results). Following each normalization, using the plotMDS function, a multidimensional scaling plot was generated from 500 loci exhibiting the highest variation in expression level among samples. Differences in transcriptome size were approximated using the sum of normalized read counts per cell; however, Limma-Voom normalizations use log-counts, which are not applicable to straight summation, so we used the DEseq package (Anders and Huber, 2010) to compute per-cell normalized counts for this purpose only. Next, a differential expression analysis was run using the normalization approaches above described, with loci identified as differentially expressed (DE) using a 0.05 p-value, 0.05 false discovery rate, and cutoff of 1 log-fold change (logFC). Differentially expressed loci were binned both broadly across the three normalization methods and more finely to characterize interplay between the three normalization results using gplots (Warnes *et al*., 2009). Gene ontology (GO) terms were extracted from the Trinotate annotations of the mapping reference. Each of the fine-scale bins of DEGs was tested for functional enrichment using GOSeq (Young *et al*., 2010).

## Results

*Quantifying cell size and density* -- The mean diploid and tetraploid 1C DNA concentrations per leaf punch extraction were 1.32591 +/- 0.09439 ug/ml and 0.76432 +/- 0.0619 ug/ml, respectively; these differed significantly (p < 0.0001) (Figure 3A). The mean tetraploid 1C DNA per punch was 57.6% of the diploid value. Following tissue digestion, the estimated values of mean number of cells per leaf punch in diploids and tetraploids were 412,146 +/- 38,002 and 268,958 +/- 20,522, respectively. These values differed significantly (p = 0.0129). The tetraploids on average had 65.3% as many cells per leaf punch as found in the diploids (Figure 3B).

**Figure 3:**
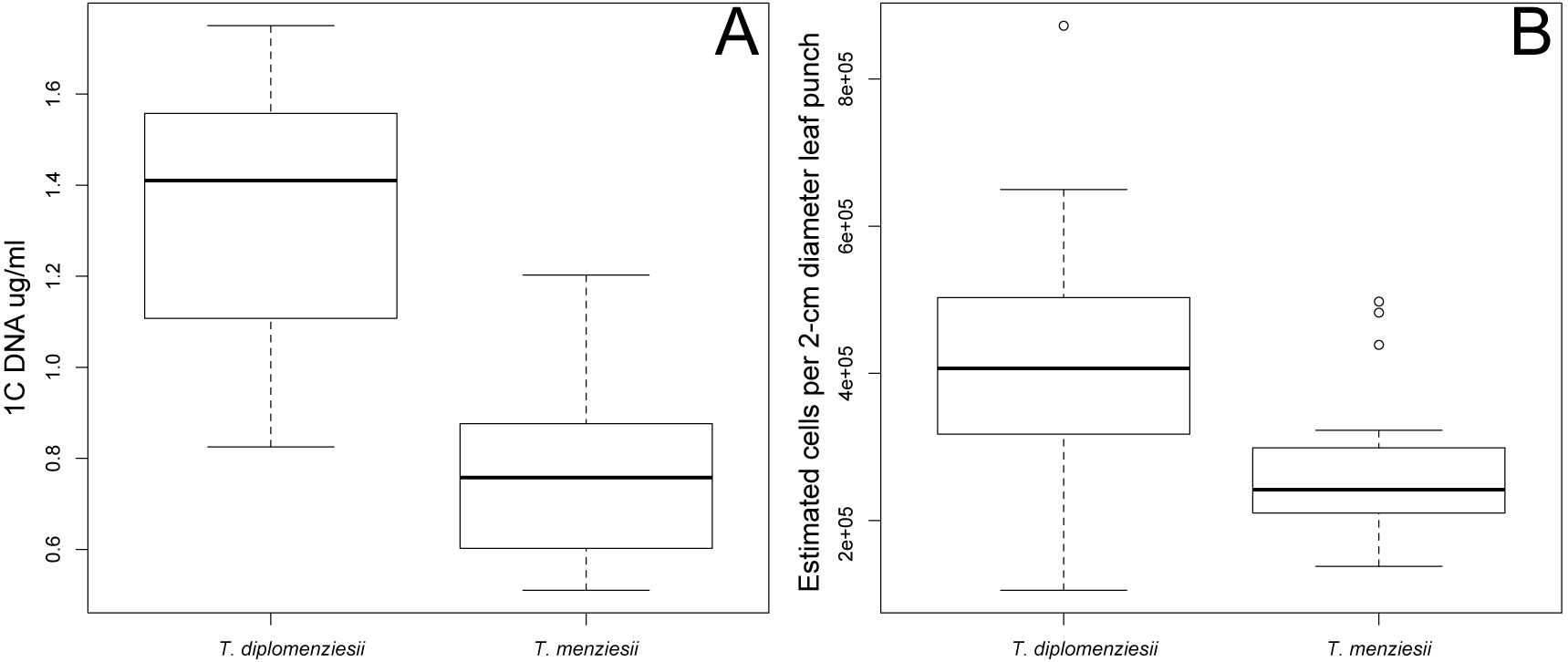
Results of ploidy variation in leaf cell density using, A: 1C DNA concentration following a DNA/RNA co-extraction, and B: estimated cell counts per 2cm diameter leaf punch.

Our two methods for characterizing differences in cell density both revealed similar reductions in tetraploid cell density; tetraploids possessed ~58% and 65% of the diploid density, based on the DNA/RNA co-extraction and cell-count results, respectively. We elected to normalize our per-cell analysis by applying a 0.65 factor to the per-cell normalization factor of the tetraploid samples to reflect the more conservative estimate of differences in cell density; however, both estimates of cell density reveal the same the major conclusions of this study (the alternative cell density difference can be run using the supplemental ipython notebook – Appendix 1).

*Differential expression analysis*—We obtained an average of 25 million reads per sample after cleaning low-quality reads (see appendix 2 for total reads per sample). The *Tolmiea* reference transcriptome assembly resulted in 58,046 isoforms binned within 28,467 clusters (henceforth genes) with an N50 of 1,821 bp. After read mapping, 26,816 genes had at least 5 non-zero counts and were used for downstream differential expression analyses. In all, 15,205 genes were annotated according to GO using Trinotate.

A multidimensional scaling plot of the 500 loci with the highest variation in expression level revealed that all three normalization methods (per transcriptome, per cell, and per biomass) performed well at clustering members from the same population with one another (Figure 4A-C). After summing the read counts normalized per cell for each sample, we found that the mean transcription per cell (henceforth transcriptome size) for the tetraploid was 2.1 times higher than the mean for the diploid (Figure 5). Additionally, the total transcriptome size per cell was highly variable, more so in the tetraploids than the diploids (20,815,579 and 11,799,791 normalized counts, respectively -- see appendix 1 for standard deviation within each population). Across the three normalization methods, the differential expression analysis found the tetraploid relative to the diploid had 1,559 up- and 1,071 down-regulated genes per transcriptome, 1,440 up- and 1,550 down-regulated genes per biomass, and 3,005 up- and 751 down-regulated genes per cell (Figure 4D-F). Across the three different normalization methods, we found 4,555 unique loci were DE under one or more methods (Figure 6). Finer binning of the interactions between normalization methods revealed the majority of DEGs to be either up-regulated in the tetraploid across all normalizations (1,392 – Figure 7A) or only up-regulated per cell (1,398 Figure 7D).

**Figure 4:**
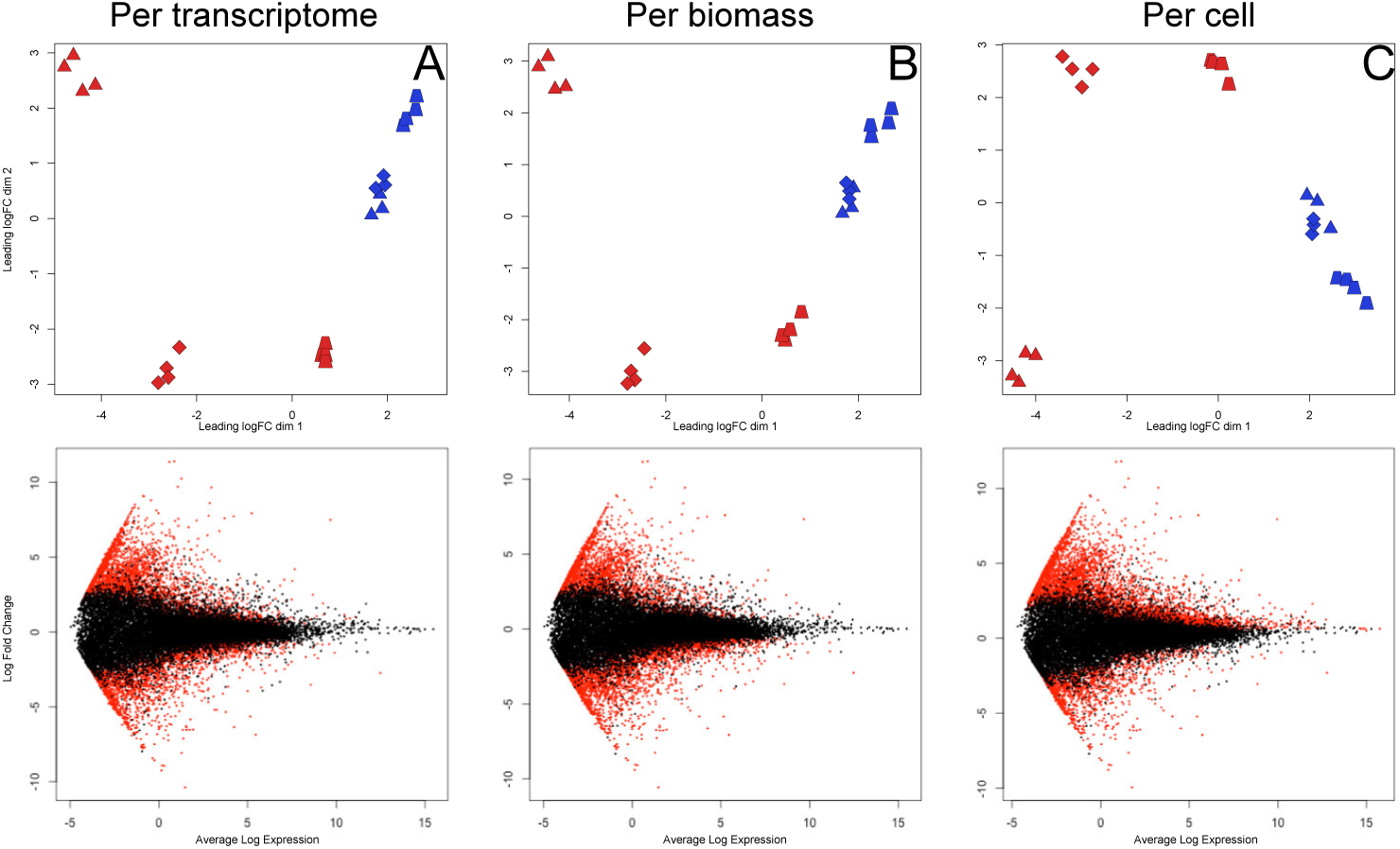
Results from multiple differential expression analysis. MDS plots A-C cluster individual based on the 500 most variable loci, with color indicating ploidal level and shape reflecting population of origin. MA plots D-F show every locus in the *Tolmiea* transcriptome (represented as dots), with log fold expression level differences in the polyploid relative to the diploid on the y-axis and average expression level on the x-axis-- red indicates statistical significance.

**Figure 5:**
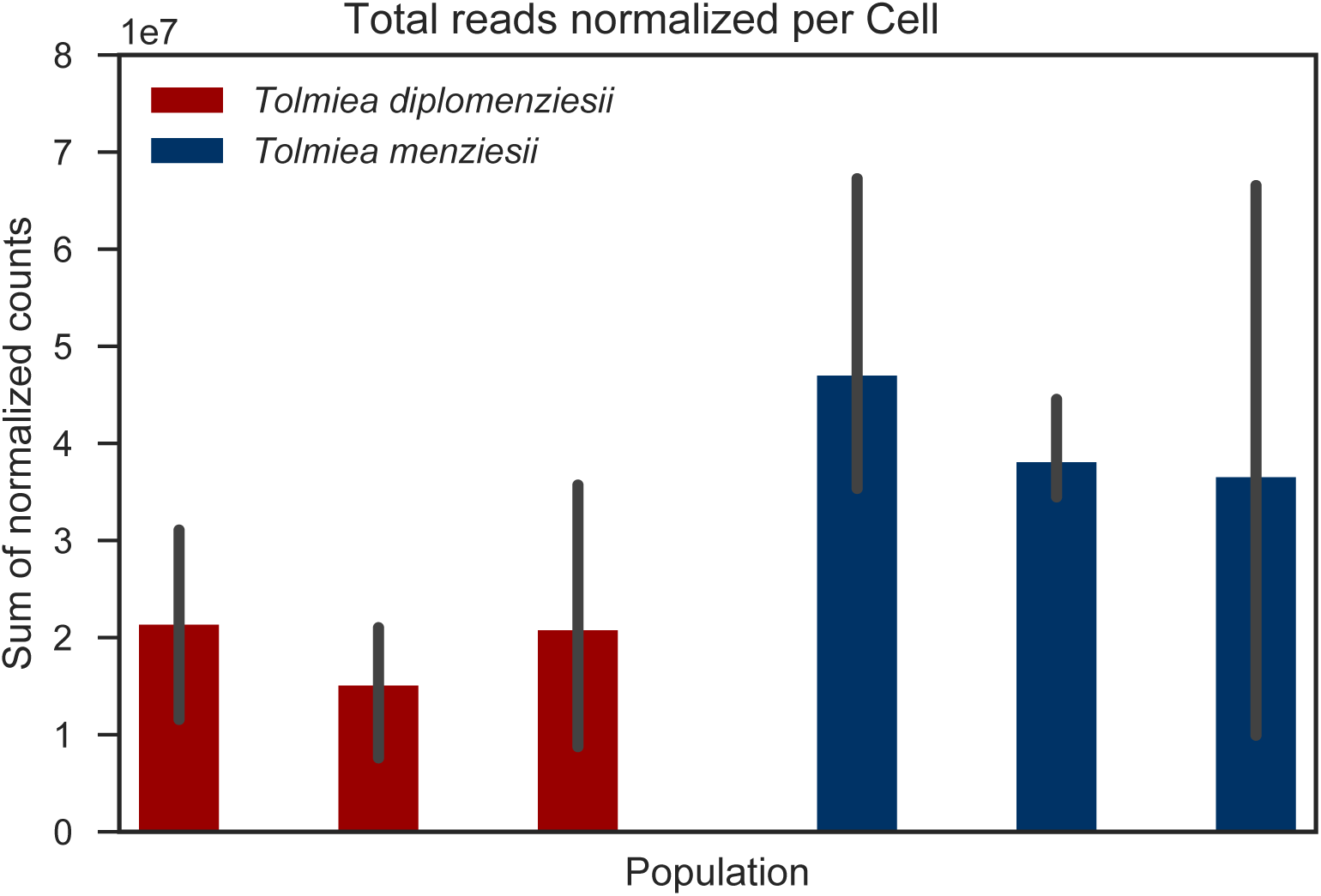
Sum of read counts normalized per cell, and clustered by population of origin. Diploid and tetraploid mean significantly differed (p < 0.008).

**Figure 6:**
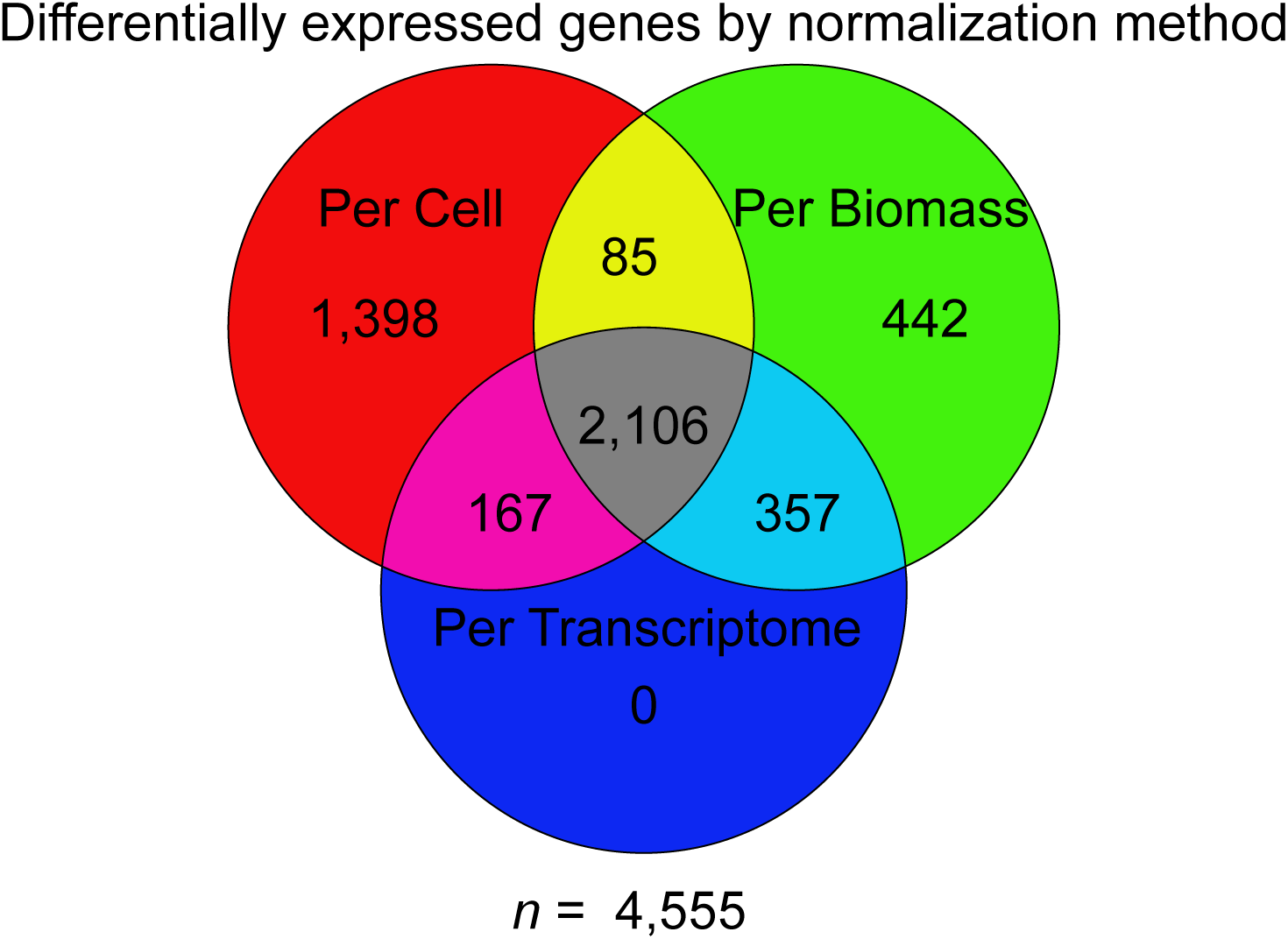
Venn diagram contrasting the three normalization methods. Numbers within the different sections indicate loci that were identified as being differentially expressed between *Tolmiea menziesii* and *T. diplomenziesii*.

**Figure 7:**
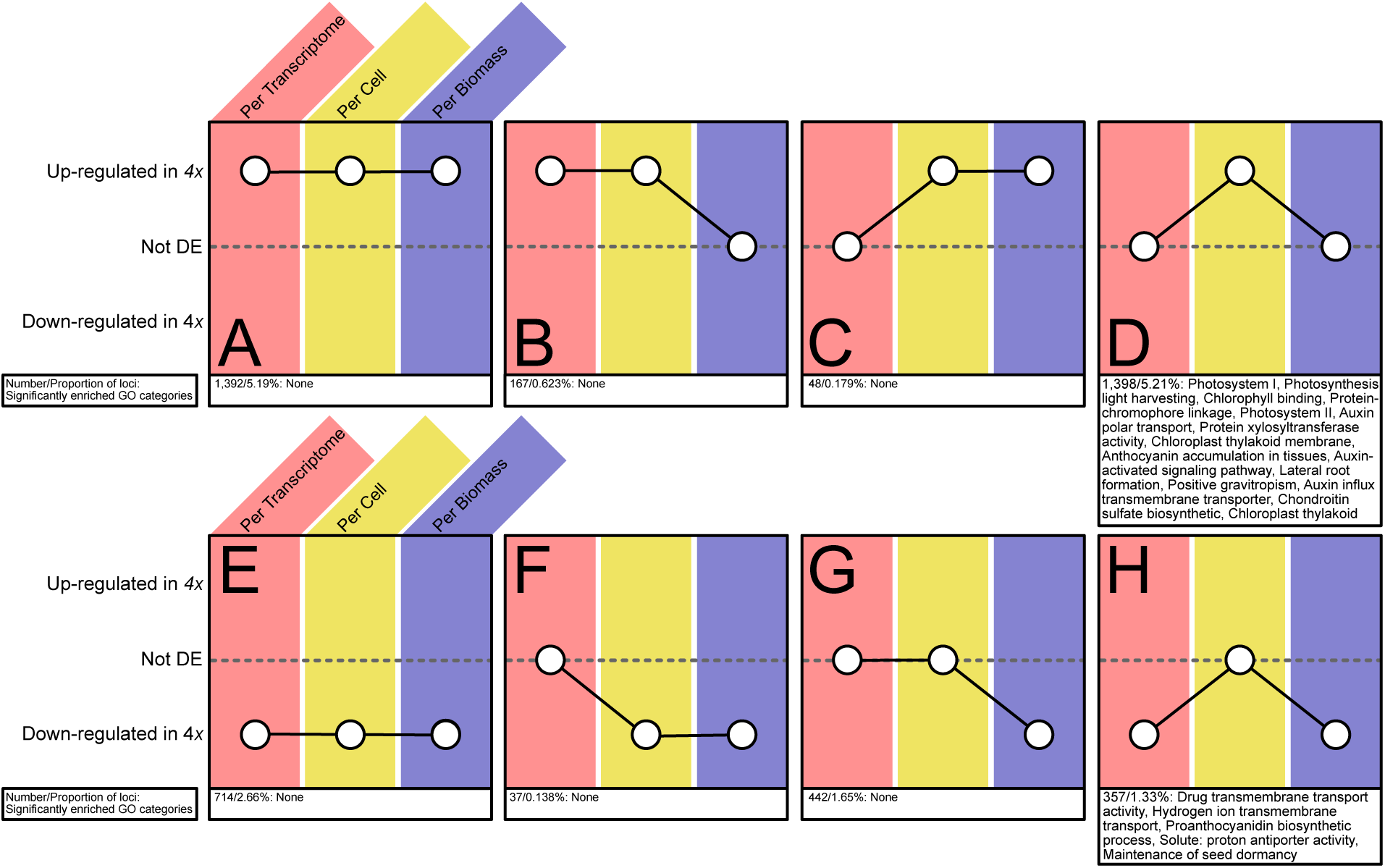
Loci binned by their DE categorization across the three normalization approaches. The number of loci belonging to each bin and the results of GO enrichment analyses are reported below the corresponding bin. Bins containing no loci are not shown.

## Discussion

### Differences in gene expression following autopolyploidy

This is the first study to leverage synthetic RNA standards to characterize differences in gene expression level per transcriptome, per biomass, and on a relative per-cell basis between ploidal levels. Through the use of this ‘three normalization approach’ and characterization of gene expression levels in diploid and autotetraploid *Tolmiea*, we found 4,555 out of 26,816 loci were DE (~17% of the transcriptome) and identified four notable trends. First, the per-transcriptome normalization, the method researchers typically use, captured the fewest DEGs and failed to detect any DEGs not found by the other two methods. Second, most differential expression occurs on a per-cell basis, and there is a clear unbalanced distribution of up- versus down-regulation in the autotetraploid relative to the diploid, with 3,005 up- versus 751 down-regulated DEGs per cell. Third, in the tetraploid, transcripts up-regulated per cell appear to compensate for a decreased cell density, resulting in a conservation of expression level per biomass relative to the diploid (see Figure 8). Loci exhibiting conservation per biomass were significantly enriched for functions related to photosynthesis and the chloroplast. Fourth, transcriptome size varied substantially across our dataset, with a significant increase in the inferred transcriptome size of the tetraploid relative to the diploid (Figure 5). Below we discuss each of these four trends in greater detail.

**Figure 8:**
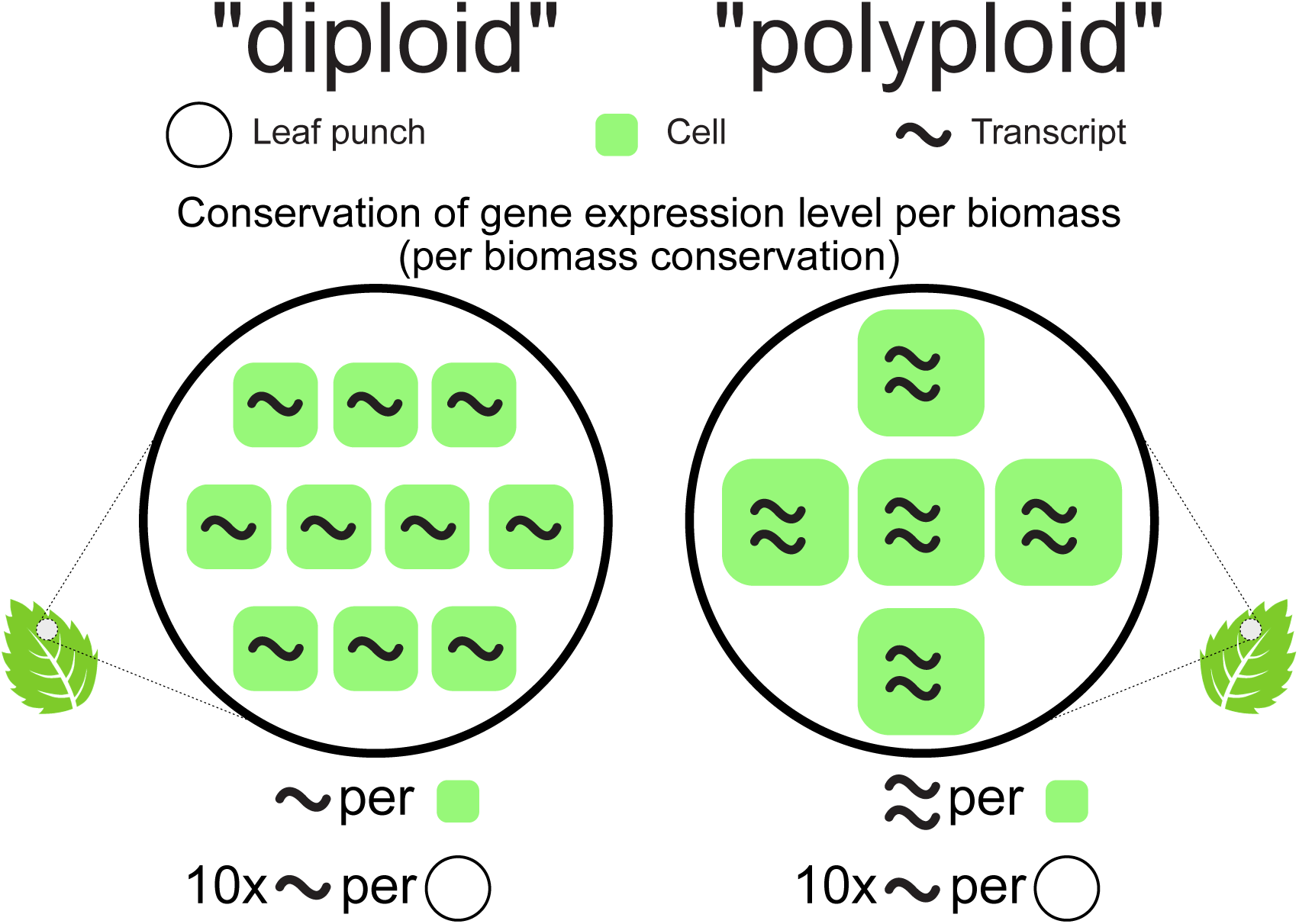
A simplified example of an interaction between expression level and cell density. =Conservation of gene expression per biomass occurs when expression level per cell in samples with lower cell density is up-regulated enough to yield equivalent levels of transcript per unit biomass.

Autopolyploids have rarely been compared to their diploid progenitors with respect to divergence in expression level (e.g., Stupar *et al*., 2007; Del Pozo and Ramirez-Parra, 2014; Zhang *et al*., 2014). These previous diploid-autopolyploid comparisons found that autopolyploids tend to deviate from diploid-like gene expression levels across 1-10% of the transcriptome (~6% in *Paulownia fortunei* – Zhang *et al*., 2014; ~10% in *Solanum phureja* – Stupar *et al*., 2007; ~1-4% in *Arabidopsis* – Del Pozo and Ramirez-Parra, 2014). However, none of these comparisons was normalized using spike-in standards; instead they used what is referred to here as a per-transcriptome comparison. Unlike the spike-in-derived per-biomass and per-cell normalized transcript counts, the results of per-transcriptome normalizations reflect concentration differences and are not a proxy for absolute expression. Therefore, the results of previous autopolyploid expression studies must be interpreted as differences in transcript concentration rather than absolute abundance. Spike-in standards have rarely been used in any evolutionary comparisons and have primarily been adopted for use in studies of model fish (zebrafish), mammals (human, mouse, and hamster), and plants (*Arabidopsis*) (e.g., Brennecke *et al*., 2013; Fomina-yadlin *et al*., 2014; Xu *et al*., 2014; Schall *et al*., 2017). Despite the use of spike-ins in these investigations of model systems, in methods development, and in single-cell sequencing, spike-in approaches have not yet been applied to evolutionarily motivated studies of natural populations. The lack of spike-in usage in non-model studies is surprising, because a reference genome is not a requirement for a spike-in-based normalization. In fact, spike-ins may be more important in non-model versus model systems because comparisons within non-model systems are often broad (e.g., interspecific comparisons) and may be more likely to result in unequal transcriptome size between treatment groups.

Considering only the results of our per-transcriptome normalization, approximately 9% of the *Tolmiea* transcriptome was differentially expressed on a concentration basis between the diploid and autotetraploid species, in line with the results for other diploid-autopolyploid pairs cited above. This finding suggests that, in general, only a small fraction of the transcriptome, less than 10%, responds to autopolyploidization through novel alterations to transcript concentration. Despite differences in transcript abundance per transcriptome representing less than 10% of all loci, concentration-based differences could have important consequences for the stoichiometry of gene expression pathways. Unfortunately, although *Tolmiea* is a good evolutionary model, it is not a genetic model, and the ability to investigate specific pathways and make inferences regarding physiological impact is severely hampered. However, it would be important for future studies to test whether members of a given pathway respond similarly with respect to the maintenance of transcript abundances relative to the transcriptome, cell, or biomass.

It is also notable that although ~9% of the loci examined in *Tolmiea* were differentially expressed per transcriptome, none of these DEGs were uniquely recovered only from the per-transcriptome analysis. The majority of per-transcriptome DEGs (2,106 of the 2,630) were differentially expressed at a sufficiently high magnitude to be detected by all three normalization methods. These results suggest that the loci typically identified as DE in previous studies of polyploid gene expression represent only differences in expression level extreme enough to be detected through a significantly altered concentration. Therefore, examination of more subtle differences in expression level requires the use of normalization approaches that allow for quantitative comparisons of transcript abundance.

When we compared expression level per cell, we found that ~14% (3,756 loci) of the transcriptome was maintained in different abundances per cell between diploid and tetraploid *Tolmiea*. Importantly, the direction of differences in expression level in *T. menziesii* relative to the diploid was extremely unbalanced. Nearly all per-cell DEGs were up-regulated in *T. menziesii* (3,005 loci), with only 751 down-regulated. In addition, 1,398 of the 3,005 per-cell DEGs up-regulated in the tetraploid were unique to the per-cell normalization and not recovered under either per-transcriptome or per-biomass normalizations (Figure 7D). Taken together, it appears that most differential expression in *Tolmiea* represents a pattern consistent with dosage sensitivity, with increased levels of gene expression in the tetraploid correlating with an increase in allele copy number. This result would not have been detected under a typical library-size normalization method, as the expression of these loci is not significantly altered in concentration relative to the whole.

The over-abundance of up-regulation per cell in the tetraploid may serve as a mechanism to compensate for reduced cell density. In other words, there is tendency in *T. menziesii* for conserved gene expression levels per biomass through novelty at the cellular level (henceforth per-biomass conservation) (Figure 8). Approximately 1,398 loci, or 5.2% of all loci, in *T. menziesii* exhibit per-biomass conservation (Figure 7D). This buffering effect is achieved by per-cell up-regulation in *T. menziesii* mirroring the decrease in cell density relative to *T. diplomenziesii*. Fifteen functional categories were significantly over-represented among the loci exhibiting per-biomass conservation. Of these, seven were related to either the chloroplast or photosynthesis. Whether there is selective pressure to conserve expression per biomass is unclear, but alteration of photosynthesis either per cell or per biomass appears to be a recurring theme across diploid/polyploid comparisons (e.g., Warner and Edwards, 1993; Vyas *et al*., 2007; Coate *et al*., 2013). For example, Warner and Edwards (1993) revealed photosynthetic conservation per biomass in *M. sativa*, but an overall increase in photosynthesis per biomass following polyploidy in *A. confertifolia*. A clear trend in the effects of polyploidization on photosynthesis per biomass has yet to emerge, and like many aspects of polyploidy, it may be lineage-specific and/or require additional study systems (Soltis *et al*., 2016).

Diploid and tetraploid *Tolmiea* occur under similar light regimes in nature (Visger *et al*., 2016), and previously collected physiological data revealed no significant difference in photosynthetic rate per leaf area (Visger *et al*., 2016) (Figure 9). In *Tolmiea*, conservation of expression level per biomass may be a mechanism for the maintenance of optimal photosynthesis per biomass, facilitating the ecological conservation of light preference in *Tolmiea*. To determine if photosynthesis-related functional enrichment of conservation per biomass is indeed a key underlying molecular mechanism for buffering cell density as it pertains to photosynthesis, additional autopolyploid systems should be similarly studied. Revisiting the work of Warner and Edwards (1993) using our spike-in standard-based gene expression approach should also show a similar conservation of gene expression per biomass in polyploid *M. sativa*. Conversely, in *A. confertifolia* where polyploidy increases photosynthesis per leaf area, we might expect photosynthesis-related gene expression per cell to be increased by a factor greater than the difference in cell density between diploid and polyploid plants (Warner and Edwards, 1993).

**Figure 9:**
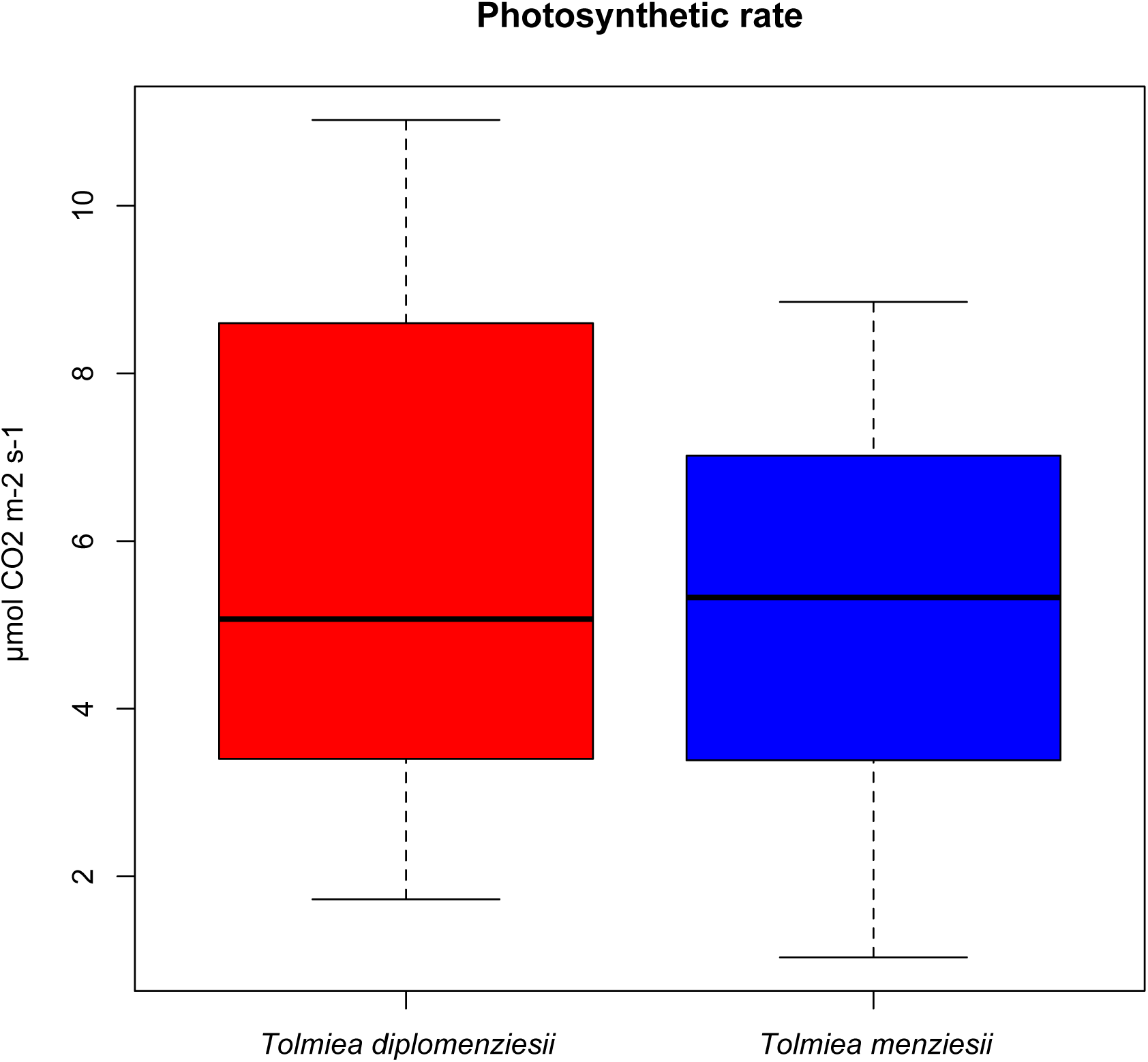
Adapted from data collected from Visger et al. 2016. *Tolmiea diplomenziesii* and *T. menziesii* did not significantly differ in photosynthetic output under common garden conditions in the green houses of University of Florida.

An initial motivation for utilizing a spike-in approach to read-count normalization was to tease apart variation in library size from differences in transcriptome size between diploid and tetraploid *Tolmiea*. We found that when normalizing read counts per cell, the mean transcriptome size of the tetraploid is over twice that of the diploid (Figure 5). This difference in transcriptome size should be qualified, as the variation within and between populations is quite large, though the mean transcriptome size still significantly differed with ploidal level. Notably, we also observed that the variability of transcriptome size was greater in the tetraploids. The diploid populations all exhibited a similar degree of variation in transcriptome size, while tetraploid populations represented both the least and greatest variation in transcriptome size (Figure 5; see Appendix 1 for standard deviation by population). Excluding all other results presented in this study, the variability of transcriptome size alone should be motivation enough for researchers of polyploidy to adopt a spike-in-based approach.

### The benefits of spike-in RNA standards

This study is not the first to apply synthetic RNA spike-in standards in dealing with a transcriptome-wide effect (e.g., see Lovén *et al*., 2012; Fomina-yadlin *et al*., 2014). However, this is the first study to: 1) leverage spike-in standards in a cross-ploidy comparison, and 2) quantify expression level on three different, biologically relevant scales. By using multiple read-count normalizations, with and without spike-in standards, we investigated the interaction of expression level per cell and per biomass between diploid and autotetraploid *Tolmiea*. Had this study been performed without the use of spike-in standards, the results would have been limited to those of the per-transcriptome normalization.

Spike-in normalization is most valuable when evaluating differences in gene expression levels between two groups that have differing cell density and/or transcriptome size. By normalizing RNAseq count data by the abundance of spike-in standards, sequencing depth and transcriptome size are effectively disentangled. Methods employed by previous comparative studies of diploid/polyploid pairs were limited to quantifying transcriptional differences on a concentration-basis only. Concentration-based comparisons, while useful for inferring alterations of pathway stoichiometry, are effectively blind to large proportions of the transcriptome exhibiting additive expression levels associated with polyploidy. That is, if the expression of many genes is increased in a single direction and is commensurate with the increase in ploidal level, then the impact on any single gene’s concentration will be minimal. An additional advantage of employing a spike-in normalization is that transcript abundance can be independently quantified relative to biomass and relative to cell. Spike-in reads can also be removed for some downstream analyses, allowing for a typical library-size normalization so that concentration-based differences may be characterized as well.

The three normalization approaches presented here are all individually informative, and the decision to include any or all of them should be guided by the research question. For example, if the research question revolves around the bulk production of a compound, evaluating differences in expression level per biomass may be the best approach. Research questions focusing on complex gene pathways may be better served by an analysis of expression level per cell. Information on potential differences in expression level stoichiometry can be gained using traditional comparisons per transcriptome. Additionally, as demonstrated by our study of *Tolmiea*, the interaction among multiple normalization approaches can be even more informative than any single approach.

While the use of non-concentration-based normalizations can enable researchers to address new questions, there is an important caveat that could lead to potential biases or increased uncertainty. Differences in RNA extraction efficiency between treatment groups are difficult to tease apart from variation in total transcriptome size. In much the same way that variation in transcriptome size influences estimates of expression level, differences in extraction efficiency could bias calculations of expression level for per-cell and per-biomass analyses. For example, if RNA extraction were half as efficient in one treatment group versus another, then differential expression analyses per cell would only consider half of the actual transcript abundance of one group relative to the other. Future approaches should consider partially accounting for this issue through the addition of a second unique set of spike-ins prior to RNA extraction. Comparing the ratio of pre-versus post-extraction spike-in abundance should highlight differences in extraction efficiency. However, even the use of a second spike-in set will not account for different extraction efficiencies if those differences arise from variation in cell lysis.

In summary, this study has demonstrated that the use of synthetic RNA spike-in standards can be used to explore previously uninvestigated aspects of divergence in gene expression levels in a comparison of a diploid and its autotetraploid derivative. To our knowledge, this multiple normalization approach has recovered the largest fraction of a transcriptome as DE in a diploid/autopolyploid species pair ever reported (~17% in *Tolmiea* vs. up to ~10% in several other plant systems; Stupar *et al*., 2007; Del Pozo and Ramirez-Parra, 2014; Zhang *et al*., 2014). Further, our methodology allowed for a fine-scale examination of how divergence in gene expression level interacts with cell density, revealing a mosaic of difference in transcript concentration, abundance per cell, and abundance across tissue.

While we compared a diploid-autopolyploid pair, in which a difference in transcriptome size might be predicted, variation in transcriptome size is rarely investigated and could be widespread in biological systems at diverse scales due to factors other than shifts in ploidy. To date, nearly all studies of global gene expression have used normalization methods that implicitly assume transcriptome size is invariable, yet this assumption is not empirically supported. Examples of studies where variation in transcriptome size might be likely include (but are by no means limited to) comparisons between related species, between different developmental stages, and across stress treatments. Yet even in these cases, the potential for variation in transcriptome size remains ignored and uninvestigated. If transcriptome size is in fact invariable between two experimental treatment groups, then following our proposed methodology, the results of the per-transcriptome and per-cell comparisons should be identical. It is of critical importance that researchers making comparisons using RNAseq data, particularly in the broad suite of examples noted above, avoid making the assumption that transcriptome size is invariable by contrasting multiple normalization approaches.

## Acknowledgements

We are thankful to J.E. Coate for insightful discussions on normalization. Clayton J. Visger was supported by NSF Graduate Research Fellowship DGE-1315138. This research was supported in part by an NSF Doctoral Dissertation Improvement Grant DEB-1501803 and by the 1KP project.

## Author’s contributions

C.J.V., G.K.W., P.S.S., and D.E.S. planned and designed the research. C.J.V. and Y.Z. performed the experiments. C.J.V. analyzed the data. C.J.V., P.S.S., and D.E.S. wrote the manuscript.

